# Can the Insect Path Integration Memory be a Bump Attractor?

**DOI:** 10.1101/2022.04.05.487126

**Authors:** Ioannis Pisokas, Nina Kudryashova, Matthias H. Hennig

## Abstract

Many animal species are able to return to their nest after a foraging excursion without using familiar visual cues to guide them. They accomplish this by using a navigation competence known as path integration, which is vital in environments that do not have prominent visual features. To perform path integration, an animal maintains a running estimate of the distance and direction to its origin as it moves. This distance and direction estimate needs to be maintained in memory until the animal uses it to return to its nest. However, the neural substrate of this memory remains uncertain. A common hypothesis is that the information is maintained in a bump attractor’s state. We test the bump attractor hypothesis and find that its predictions are inconsistent with the path integration behaviour of ants, thus highlighting the need for alternative models of path integration memory.

## Introduction

When a foraging ant of the species *Cataglyphis fortis*, inhabiting the Saharan desert, embarks in search of food, it typically follows a circuitous path. However, once it finds a food item, it readily returns to its nest, travelling along a straight path, even though there are no visual cues to guide it along its trip (***Müller and Wehner, 1988***; ***Collett, 2019***; ***Menzel and Muller, 1996***). The animal achieves this by maintaining a running estimate of the direction and distance to a starting location, using a navigational competence known as ‘path integration’ or ‘dead-reckoning’ (***Darwin, 1873***; ***von Frisch, 1967***; ***Mittelstaedt, 1985***; ***Müller and Wehner, 1988***). This estimate is known as the ‘home vector’ (***Collett, 2019***) and is updated continuously as the animal moves (***Müller and Wehner, 1988***; ***Menzel and Muller, 1996***; ***Heinze et al., 2018***; ***Collett, 2019***).

To update its home vector, the insect has to have access to two pieces of information at every instant in time, its current heading and an estimate of the distance^1^ it travels. These displacement estimates are accumulated continuously updating the home vector as the animal travels (***Heinze et al., 2018***; ***Collett, 2019***). This procedure is similar to that employed by sea navigators in the past who used a magnetic compass to gauge their heading and vector addition to track their position on a map.

Path integration has been described in several species, including desert ants of the genus *Cataglyphis*, the honey bee *Apis mellifera*, the sweat bee *Megalopta genalis*, the fruit fly *Drosophila melanogaster*, as well as in rodents (***Müller and Wehner, 1988***; ***Collett, 2019***; ***Menzel and Muller, 1996***; ***Heinze and Homberg, 2007***; ***Heinze et al., 2013***; ***Stone et al., 2017***; ***Kim and Dickinson, 2017***; ***McNaughton et al., 2006***). The spatial scale of path integration varies between species. For instance, fruit flies (*Drosophila melanogaster*) employ path integration for returning to a previously visited drop of sugar a few centimetres away (***Kim and Dickinson, 2017***), while other insect species use path integration to return to their nest over much larger distances, for instance, hundreds of meters in the case of the desert ant *Cataglyphis fortis*, or several kilometres in the case of the honey bee *Apis mellifera* (***Sommer and Wehner, 2004***; ***Cheng et al., 2005***; ***Huber and Knaden, 2015***; ***von Frisch, 1967***).

At the end of its excursion, a path integrating animal can return to its nest by travelling in the direction and for the distance indicated by its accumulated home vector memory (***Müller and Wehner, 1988***; ***Menzel and Muller, 1996***; ***Collett, 2019***). To do this, the animal needs to be able to maintain its home vector in memory for the duration of its excursion, even if it is interrupted and prohibited from returning to its nest for several hours (***Ziegler and Wehner, 1997***; ***Cheng et al., 2005***). Therefore, a memory mechanism that can be updated quickly and maintain its state long enough is required (***Pisokas et al., 2022***). However, the substrate of the employed memory remains unknown.

Most path integration models leave the memory substrate unspecified, while some authors have suggested that the home vector memory might be maintained through reverberating neural activity and more specifically, in a bump attractor network (***McNaughton et al., 1996***; ***Samsonovich and McNaughton, 1997***; ***Conklin and Eliasmith, 2005***; ***McNaughton et al., 2006***; ***Burak and Fiete, 2009***; ***Vickerstaff and Di Paolo, 2005***; ***Haferlach et al., 2007***; ***Kim and Lee, 2011***; ***Goldschmidt et al., 2015***; ***Webb and Wystrach, 2016***; ***Stone et al., 2017***; ***Goldschmidt et al., 2017***).

Bump attractors are recurrent neural networks that maintain a localised peak of neural activity. A bump attractor can be used to encode an agent’s spatial position as the location of the peak of neural activity (‘activity bump’) in the network (***Figure 1A***). If we imagine that the neurons of the bump attractor are organised topologically (if not spatially) on a line, the higher the represented spatial value, the further along the line the most active neurons will be located (***Figure 1A***). This constitutes a positional encoding of value along a line of neurons.

**Figure 1.**
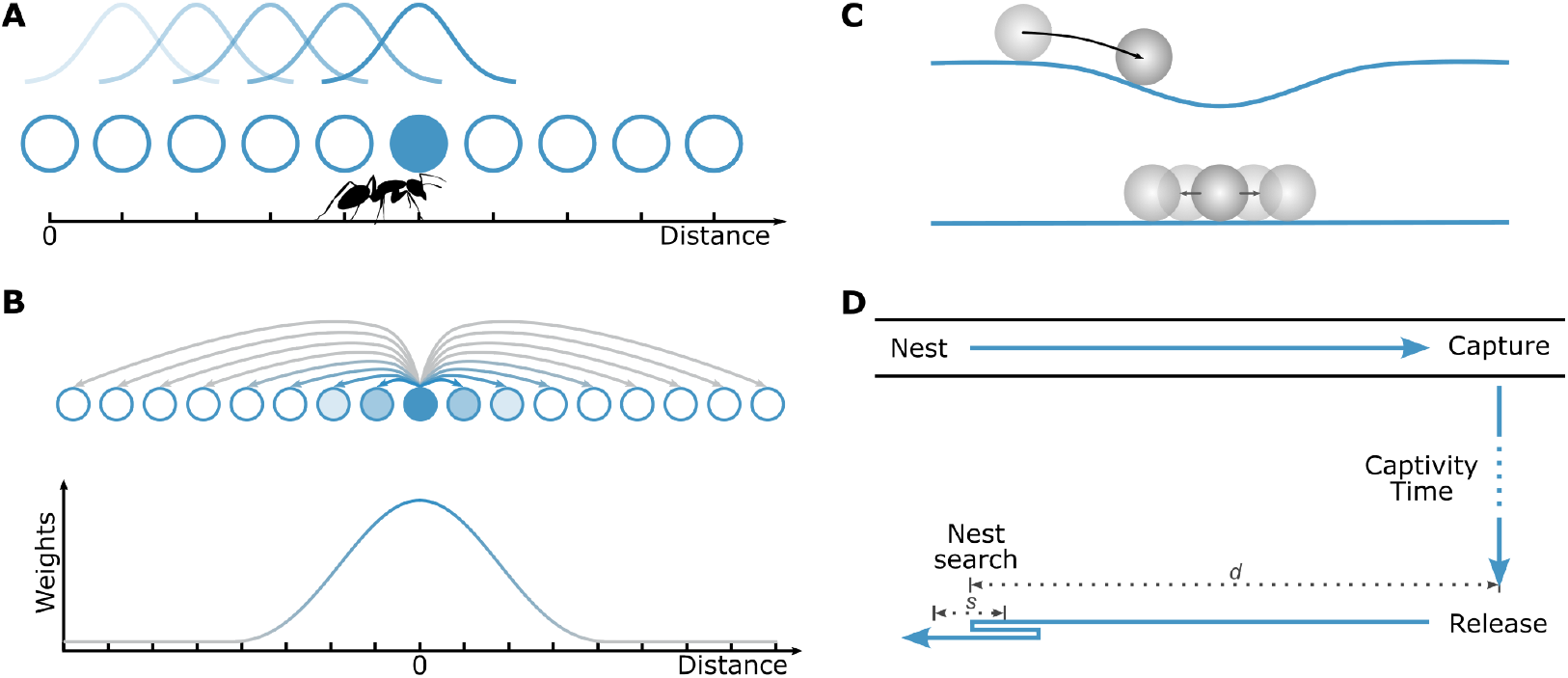
Bump attractor networks as spatial location memory. (**A**) The state of a bump attractor is encoded in the position of the most active neuron in the network. In a bump attractor that integrates an animal’s velocity, the location of the activity bump would encode the spatial position of the animal. (**B**) In a bump attractor network, the synaptic connectivity weights pattern results in the formation of one activity bump. The synaptic weights of one exemplar neuron are shown; the same synaptic pattern is repeated for all neurons. The synaptic connections (arrows) are colour-coded by their strength, with blue denoting excitatory synapses and grey inhibitory synapses. The synaptic strength is a function of the distance from the emanating neuron. (**C**) Inhomogeneities in the biophysical and synaptic properties of the network cause a systematic drift in the bump’s location (top of panel), while neuronal noise causes an unbiased stochastic drift of the bump’s location (bottom of panel). (**D**) For measuring the homing accuracy of path integration, ants were captured once they reached a feeder, held in captivity for different amounts of time, and then released in a remote unfamiliar location (***Ziegler and Wehner, 1997***). Upon release, ants that had not been kept in prolonged captivity would typically run towards their expected nest location, not finding it since they were displaced, and would perform a focused search for the entrance of their nest. The distance (*d*) at which ants start searching for their nest is a measure of the memorised home vector distance. The spread of this distance (*s*) over trials is a measure of the homing distance error or dispersion.

In bump attractor networks, the activity bump is maintained through specific synaptic connectivity with short-range excitation and long-range inhibition (***Gerstner et al., 2014***, see ***Figure 1B***). The dynamics of the neuronal activities are described by the coupled set of differential equations

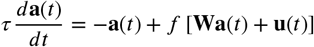

where *τ* is the neuronal time constant, **a** is the neuronal activities vector, *f*[·] is the neuronal activation function, **W** is the weight matrix stipulating a local lateral excitation profile between neurons and global inhibition (section Bump attractor network model in Materials and Methods), and **u** (*t*) is the vector of external inputs to the attractor network.

The location of the activity bump can be changed by external input that breaks the balance between excitation and inhibition to move the bump towards a particular direction. With appropriate connectivity, the amount of the bump’s movement along the neural network can be proportional to the integral of the input stimulus over time (***Skaggs et al., 1995***). In this case, bump attractor networks can be used as neural integrators that encode the integral of the input signal as the location of the activity bump along the network.

Bump attractor networks have been used to model systems that gradually accumulate velocity signals, such as angular velocity for tracking the head direction of rodents (***Zhang, 1996***; ***Redish et al., 1996***; ***Goodridge and Touretzky, 2000***; ***Boucheny et al., 2005***) or translational velocity in path integration models of rodent place and grid cells to track the spatial position of the animal (***McNaughton et al., 1996***; ***Samsonovich and McNaughton, 1997***; ***Conklin and Eliasmith, 2005***; ***Burak and Fiete, 2009***).

In insects, the computations pertaining to path integration are believed to occur in the central complex, a brain structure conserved among insect species (***Heinze et al., 2013***; ***Pfeiffer and Homberg, 2014***; ***Seelig and Jayaraman, 2015***; ***Weir and Dickinson, 2015***; ***Heinze, 2015***; ***Turner-Evans and Jayaraman, 2016***; ***el Jundi et al., 2018***; ***Franconville et al., 2018***). Neurons that encode the animal’s heading and speed have been identified in the central complex (***Heinze and Homberg, 2007***; ***Homberg et al., 2011***; ***Seelig and Jayaraman, 2015***; ***Green et al., 2017***; ***Stone et al., 2017***; ***Lyu et al., 2022***; ***Lu et al., 2022***; ***Hulse et al., 2021a***) and its characteristic columnar structure and regular projection patterns have been hypothesised to provide a plausible substrate for the required path integration computations (***Stone et al., 2017***).

The existence of the necessary anatomical connectivity for the formation of a bump attractor has been demonstrated in the head direction circuit of insects (***Seelig and Jayaraman, 2015***; ***Kim et al., 2017***; ***Turner-Evans et al., 2017***; ***Kakaria and de Bivort, 2017***; ***Green et al., 2017***; ***Su et al., 2017***; ***Green and Maimon, 2018***; ***Pisokas et al., 2020***; ***Turner-Evans et al., 2020***), but no direct connectomic evidence of bump attractor synaptic structure for encoding translational displacement has been identified in the underlying neural substrate.

Current bump attractor models exhibit limited state stability over time (***Brody et al., 2003***), while insects can maintain the memory of their home vector for hours (***Ziegler and Wehner, 1997***). This is an important discrepancy and it is, therefore, imperative to investigate whether bump attractors are an ecologically plausible underlying memory mechanism for path integration. To address this question, we compare the dynamics of bump attractors with those of the home vector memory of the desert ant *Cataglyphis fortis*. If the ants’ memory mechanism is indeed a bump attractor network, the state dynamics of such a network should be consistent with the behaviour dynamics of the ants.

## Results

### Homing accuracy

The way the homing accuracy of an animal degrades over time could provide crucial information about its spatial memory characteristics. The state of a bump attractor is subject to two phenomena: systematic and stochastic drift. The systematic drift is due to inhomogeneities in the network parameters, while the stochastic drift is due to neuronal noise. Both of these would affect the bump’s location (***Figure 1C***). In bump attractors, when no input signal is provided, stochastic neuronal noise causes the activity bump’s location to stochastically drift over time (***Compte et al., 2000***; ***Burak and Fiete, 2012***). Due to the stochastic drift, the bump increasingly deviates from its original location resulting in increased dispersion over time (***Figure 2A***). Therefore, if the ant’s home vector memory were based on a bump attractor network, we would expect an increase in the animal’s home vector memory error and accordingly a deterioration in its homing distance accuracy over time.

**Figure 2.**
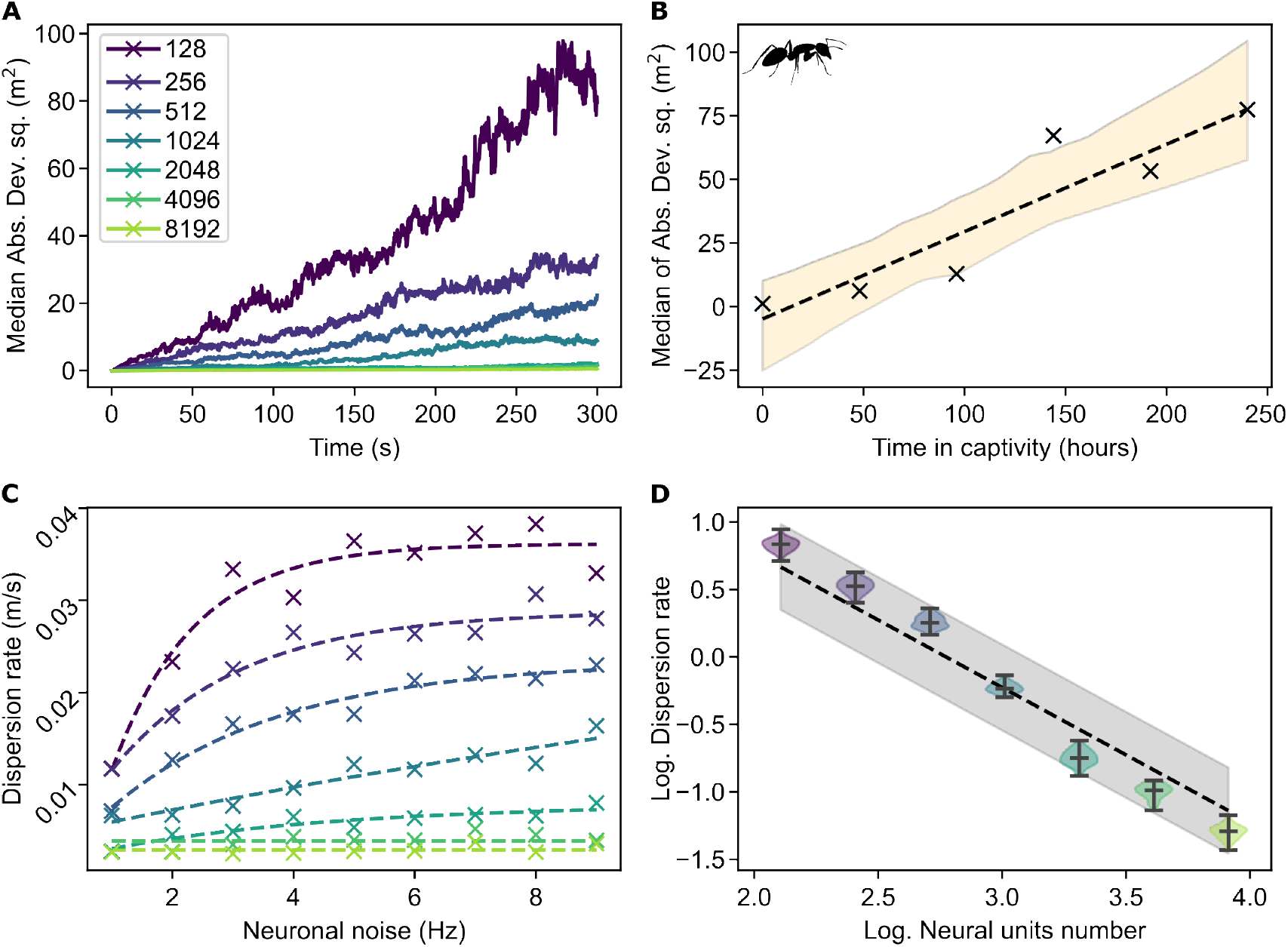
Path integration error increase over time. (**A**) Simulation results of the error (dispersion, measured as squared median absolute deviation) in the attractor’s activity bump location over time for networks of different sizes (300 trials for each condition). Since a homogeneous neuronal population was used, the error increase was due to neuronal noise. The error increases linearly with time. An increased number of neuronal units in the bump attractor network results in a reduced error increase rate. (**B**) The ants’ homing distance error increases linearly with captivity duration (Data from ***Ziegler and Wehner 1997***). The line *y* = *ax* + *b* was fitted to the data with parameters *a* = (0.343 ± 0.088) m^2^/h and *b* = (−4.784 ± 12.122) m^2^ (mean ± std, R^2^=0.85), showing that the homing distance error (dispersion) increased at a rate of 0.343 m^2^/h. The 95% confidence intervals for the linear regression fits are indicated by the shaded zone. (**C**) Error increase rate (dispersion rate, measured as median absolute deviation per second) in the activity bump’s location for different attractor network sizes. The colours signify the numbers of neurons used in the simulations (as in **A**). (**D**) The required number of neurons is inversely proportional to the dispersion rate 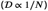. The function (dashed line) *y* = −*x* + *b* was regressed to the logarithm of the dispersion rate vs. the logarithm of the neurons number with parameter *b* = 2.77 ± 0.02 (mean ± std, R^2^=0.95). The 95% confidence intervals for the dispersion rate are indicated by the shaded zone. Extrapolation to dispersion rate 0.343 m^2^/h gives a minimal required network size of around 6 million neuronal units (6.2 × 10^6^ ± 2.3 × 10^6^ neuronal units, mean ± std).

Two studies have attempted to quantify how the homing accuracy of path integrating ants (*Cataglyphis fortis*) degrades with time (***Ziegler and Wehner, 1997***; ***Cheng et al., 2005***). The *Cataglyphis fortis* ants are endemic in the salt pans of Tunisia and Algeria, where the skyline is flat, providing no prominent visual cues the animals can use for navigation, so the ants are known to resort to path integration for their navigation (***Müller and Wehner, 1988***). Therefore, this ant species is a suitable candidate for studying path integration in isolation from other navigational competencies. In the aforementioned studies, the ants were captured once reaching a feeder, away from their nest, and were kept confined in a dark box for different amounts of time before being released at an unfamiliar location. Once released, these ants typically run towards their expected nest location (***Figure 1D***). The authors measured the distance the ants ran before they started searching for their nest’s entrance.

Even though the typical foraging excursion of a *Cataglyphis fortis* ant in its natural habitat lasts no more than one hour, ***Ziegler and Wehner*** (***1997***) found that the ants maintained their home vector memory for several hours, a finding that was confirmed by ***Cheng et al**.* (***2005***) in a similar experiment. We analysed the data reported by these authors and observed that the homing distance error (defined here as the squared median absolute deviation, a measure of dispersion) increases with a constant rate over captivity time (***Figure 2B***). In bump attractors, stochastic drift causes a constant increase in the bump’s expected location variance over time (***Compte et al., 2000***; ***Burak and Fiete, 2012***). Similarly with the variance, the squared median absolute deviation of the bump’s expected location would grow linearly with time, as seen in ***Figure 2A***. This is in agreement with the linear relationship between captivity time and squared median absolute deviation seen in the ants (***Figure 2B***). However, more aspects of the animal behaviour should coincide with the predictions of the bump attractor hypothesis for it to be a plausible memory substrate.

### Required number of neurons

In principle, a bump attractor could reproduce the linear increase of homing distance error over time (***Figure 2***A). However, the homing distance error of the *Cataglyphis fortis* ants increases at a rate of 0.343 m^2^/h (***Figure 2B***), which is three orders of magnitude slower than the bump attractor simulations in ***Figure 2***A. In bump attractor networks, the dispersion (squared median absolute deviation) of the bump’s expected location depends on the number of neurons in the network (***Figure 2A***). Thus the question arises, how many neurons are required to reproduce the dispersion rate observed in the ant behaviour experiments?

The required number of neurons, *N*, is inversely proportional to the desired dispersion rate 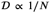 (***Compte et al., 2000***; ***Burak and Fiete, 2012***). However, the exact number of neurons depends on the assumptions about the circuit structure and the biophysical properties of the neurons. We, therefore, provide an indicative solution for a bump attractor implementation in a spiking neural network with physiologically plausible neuronal parameters (see Materials and Methods). The result should be seen as an indication of order of magnitude and not as a specific number of required neurons. We varied the size of the simulated bump attractor network and used the above relation to extrapolate from the simulation results to larger networks, estimating the size of the network required to reproduce the experimentally measured dispersion rate of the ants (***Figure 2D***, dashed line). We found that a bump attractor network of at least 6 million neuronal units is required (6.2 × 10^6^ ± 2.3 × 10^6^ neurons, mean ± std) for exhibiting a dispersion rate comparable with that observed in the ants’ behaviour (0.343 m^2^/h). In reality, each neuronal unit in a bump attractor might consist of several neurons, and the circuit would require additional neuronal resources for controlling the bump’s shift in response to input signals as well as a population of inhibitory neurons. This means that the actual number of neurons required for the bump attractor circuit would be a multiple of 6 million.

In insects, the home vector memory has been speculated to lie in the upper central body (CBU, also known as the fan-shaped body) of the central complex (***Stone et al., 2017***; ***Collett, 2019***). The exact number of neurons in the ant’s brain is not known, but the comparable central complex of *Drosophila melanogaster* is estimated to have no more than 5,000 neurons, while the whole brain is estimated to have 200,000 neurons (***Raji and Potter, 2021***; ***Hulse et al., 2021b***). Current models and neuroanatomical evidence suggest the existence of 8 or 16 independent distance integrators, one for each represented cardinal direction (***Stone et al., 2017***), and independent distance integrators might exist for path integrating based on optical flow and stride counting self-movement estimation, further increasing the required number of neurons (***Collett, 2019***). Therefore, the insect brain does not have enough neurons to accommodate a memory circuit that requires so many neurons to reproduce the homing distance error rate (dispersion rate) observed in the animals.

### Required neuronal time constant

In the previous section, we assumed that neurons have a typical membrane and synaptic time constant (see section Bump attractor network model in Materials and Methods). However, the bump’s dispersion rate also depends on the neuronal time constant and decreases as the time constant increases, 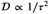 (***Burak and Fiete, 2012***, see ***Figure 3***). If we assume a network of a plausible size, i.e. a few hundreds of neurons (256 neuronal units were used in the simulations), we find that we would need a neuronal membrane time constant of around 16 h (16 h ± 8 h, mean ± std), to achieve a bump attractor with a dispersion rate of approximately 0.343 m^2^/h. This is beyond the time constants range of typical neurons.

**Figure 3.**
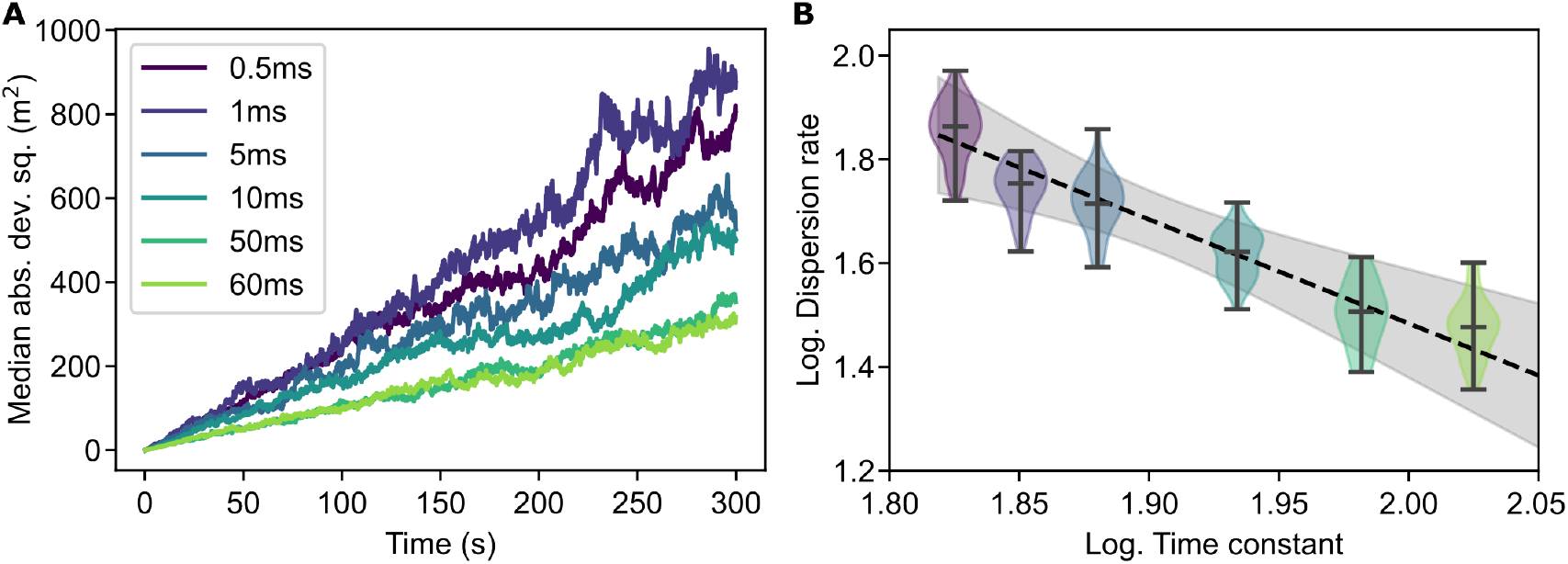
Effect of time constant on error increase. (**A**) Simulation results show the error (dispersion, measured as squared median absolute deviation) in the bump’s location over time for different neuronal membrane time constants (300 trials for each condition). Higher time constants result in lower error increase rates. A network of 256 neuronal units was used for the simulations. Note the higher error increase rate than in ***Figure 2A*** due to the use of higher Poisson neuronal noise for the attractor network to sustain a bump at higher neuronal time constants (1400 impulses/s Poisson noise). As the time constant is varied towards lower and higher values, its effect is progressively reduced until the model breaks. (**B**) Dependence of error increase rate (dispersion rate, measured as squared median absolute deviation per second) on the neuronal membrane time constant for the simulation data in **A**. The function 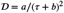 was regressed to the dispersion rate 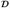 vs. the time constant *τ* simulation data with parameters *a* = (0.31 ± 0.03) m^2^ s and *b* = (66 ± 4) ms (mean ± std, R^2^=0.82). In panel **B**, the logarithm of 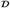 vs. the logarithm of *τ* + *b* are depicted. The shaded zone indicates the 95% confidence intervals for the model fits. Extrapolation to the desired dispersion rate, 0.343 m^2^/h (***Figure 2***B), resulted in a minimum required neuronal time constant of 16 h (16 h ± 8 h, mean ± std). **Figure 3—figure supplement 1. Bump attractor error increase using neurons with a persistent non-specific cation conductance**. Simulation results show the error in the activity bump’s location over time for networks of different sizes. A homogeneous neuronal population was used. In these experiments the dispersion of the bump’s location was significantly smaller than in ***Figure 2A*** and ***Figure 3A***.

### Neurons with a long time constant

One way to increase the time constant of neurons is to replace each neuron with a recurrent neural circuit with positive feedback (***Cannon et al., 1983***; ***Seung, 1996***). Such recurrent networks may exhibit time constants higher than those of their constituent neurons, however, at the expense of an even larger number of neurons.

On the other hand, principal neurons in layer V of the entorhinal cortex (EC) and the lateral nucleus of the lateral amygdala (LA), are able to generate graded persistent activity that is sustained at a constant frequency for prolonged periods of time (***Egorov et al., 2002***, ***2006***). These single-neuron oscillators are not dependent on reverberating activity in recurrent circuits, and their spike rate can be gradually increased or decreased with appropriate synaptic input (***Egorov et al., 2002***; ***Fransén et al., 2006***).

This graded persistent activity mechanism depends on a non-specific calcium-sensitive cationic current and introduces dynamics operating at a time scale that is many times larger than the time constants of the neuronal membrane and synapses. We, therefore, replaced the neurons in our bump attractor model with a model of these neurons to investigate whether the prolonged activity phenomenon would result in more stable bump attractor dynamics. Indeed this replacement resulted in a significantly lower bump location dispersion rate (***figure Supplement 1***), making such a bump attractor network model seem to be a plausible solution even in the face of the limited number of neurons in an insect’s brain. If such neurons exist in insect brains, the required homing distance error increase rate (rate of increase in the homing distance dispersion) of 0.343 m^2^/h could be achievable even with as few as 256 neuronal units. However, so far, graded persistent activity has only been reported in mammalian cells *in vitro* under very specific non-physiological conditions, and it is uncertain whether it is expressed under realistic physiological *in vivo* conditions.

### Homing distance decay regime

Even though we have shown that the desired dispersion rate could be achieved with neurons exhibiting persistent graded activity, there is an additional aspect of the ants’ distance memory dynamics that we should consider. In bump attractors, the bump may drift isotropically and equally likely towards higher and lower values (***Figure 4A***). Therefore, another prediction of the bump attractor hypothesis is that the increase of the animal’s homing distance error due to memory drift would be equally likely to result in longer or shorter homing distances than the actual distance to the nest. On the contrary, the homing distance of the ants systematically and monotonically decreases with time (***Figure 4B***). This is a fundamental difference between ant behaviour and the predicted state loss of bump attractors, rendering the vanilla bump attractor hypothesis an insufficient explanation of the observed animal behaviour.

**Figure 4.**
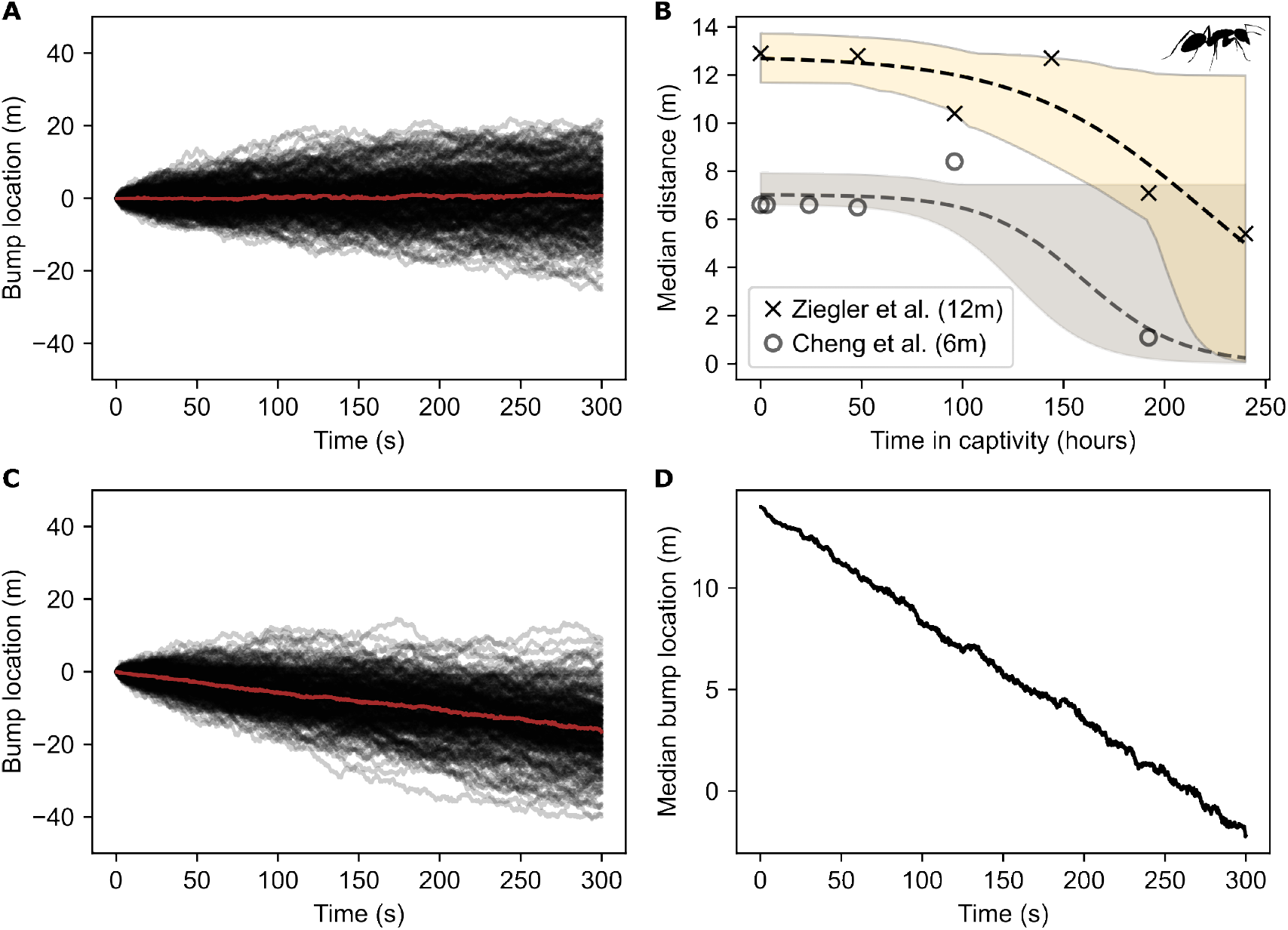
Loss of state value over time. (**A**) Dispersion in the bump attractor’s state over time for 200 simulation trials starting from the same initial state of the bump attractor. The initial bump location disperses over time due to stochastic neuronal noise. During individual trials, the bump is equally likely to move towards higher or lower values. The median across trials is shown in red. (**B**) The homing distance of ants monotonically decays with captivity time. Data points in black x’s are from ***Ziegler and Wehner*** (***1997***), while data points in grey o’s are from ***Cheng et al**.* (***2005***). The actual distance between nest and capture point is 12 m in ***Ziegler and Wehner*** (***1997***) and 6 m in ***Cheng et al**.* (***2005***). The inverse logistic function 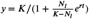 was regressed to the data points, where *t* is the captivity time in hours and the regressed parameter values are *K* = (12.800 ± 0.761) m, *N*_*l*_ = 0.099 ± 0.580, and *r* = 0.022 ± 0.035 (mean ± std, R^2^=0.85) for ***Ziegler and Wehner*** (***1997***) and *K* = (7.000 ± 0.357) m, *N*_*l*_ = 0.010 ± 0.006, and *r* = 0.041 ± 0.018 (mean ± std, R^2^=0.86) for ***Cheng et al**.* (***2005***). The shaded regions indicate the 95% confidence intervals for the function fits. (**C**) Effect of systematic anisotropy in the synaptic weights of the bump attractor. Anisotropy was introduced by changing the synaptic strength pattern from Gaussian to skewed Gaussian with parameter a=−0.0005, resulting in a constant bump drift rate. Data of 200 simulation trials are shown (the median across trials is shown in red). The bump location changes because of the combination of systematic drift due to the structural bias and stochastic drift due to neuronal noise. (**D**) Median bump location of the data shown in **C** with the starting value set to 15 m for comparison with **B**. Unlike the ants, the bump’s mean location drifts with a constant rate.

It is, however, conceivable that the monotonic decay of the homing distance over time might be due to a systematic bias in the bump attractor network. This systematic bias could be caused by a structural bias in the synaptic weights that would shift the activity bump in one direction over time. We, therefore, tested the effect of introducing bias to the synaptic weights of the bump attractor by replacing the Gaussian synaptic profile (***Figure 1B***) with a skewed Gaussian synaptic profile (see section Bump attractor network model in Materials and Methods). This manipulation resulted in systematically biased drift in the bump’s location; however, the activity bump moved at a constant rate (***Figure 4C*** and ***Figure 4D***) that cannot account for the accelerating decay rate seen in ants (***Figure 4B***). This suggests that a different process causes homing distance degradation in the animals.

Altering the synaptic bias to produce the accelerating decay regime observed in the animals would require the biased synaptic profile to change depending on the homing distance. Such dynamic adaptation of the synaptic weights seems unrealistic and would require an additional neuronal apparatus to manage it, raising the number of needed neurons once more.

### Effect of bump dispersion on agent homing

In animals, memory does not exist in isolation but in the context of behaving agents. We, therefore, investigated the bump attractor memory model as part of a simulated agent. This allowed us to compare the agent’s behaviour with that of the animals and investigate alternative memory degradation models. After travelling away from its origin (nest), the agent was held in ‘captivity’ for different waiting times, and then released for returning to its nest.

Theoretical analysis, neuroanatomical evidence, and modelling work indicate that the insect home vector is stored as a Cartesian vectorial representation with individual memory units storing the coordinate values along each cardinal axis (see ***Figure 5***; ***Cheung and Vickerstaff, 2010***; ***Vickerstaff and Cheung, 2010***; ***Wittmann and Schwegler, 1995***; ***Kim and Hallam, 2000***; ***Vickerstaff and Di Paolo, 2005***; ***Haferlach et al., 2007***; ***Stone et al., 2017***; ***Wolff and Rubin, 2018***; ***Hulse and Jayaraman, 2020***; ***Pisokas et al., 2022***). Modelling has shown that the columnar organisation of the central complex provides a neural basis for potentially encoding and storing the coordinate values along eight columns of neurons, with each column corresponding to a cardinal axis (***Haferlach et al., 2007***; ***Stone et al., 2017***). In this model, the eight memory values form a sinusoidal pattern with its amplitude encoding the distance the agent has travelled away from its origin and the location of the minimal memory value (column) corresponding to the direction the agent has mostly travelled away from its origin (***Figure 5C***) (***Hartmann and Wehner, 1995***; ***Haferlach et al., 2007***; ***Stone et al., 2017***). When a homing agent returns to its origin, the sinusoid’s amplitude approaches zero and does not provide the agent with a direction to move towards. As a result, the homing agent moves in loops around the current location, resembling the search pattern observed in ants (***Stone et al., 2017***).

**Figure 5.**
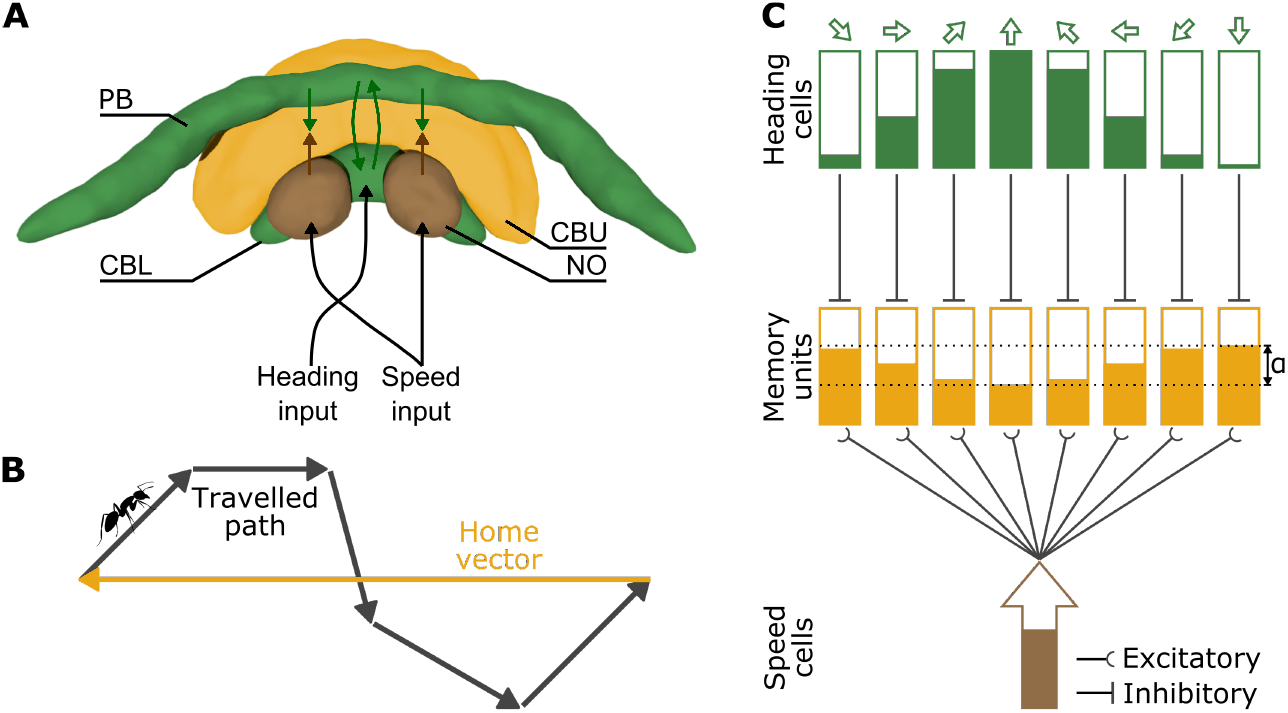
A Cartesian home vector encoding model of insect path integration. (**A**) The anatomy of the central complex and the main neuronal signal pathways involved in path integration. The brain structures — protocerebral bridge (PB), upper central body (CBU, also fan-shaped body), lower central body (CBL, also ellipsoid body), and noduli (NO) — are colour coded to match the conceptual drawing in **C**. The compartment and arrow colours indicate the type of identified signals (heading and speed) and their convergence in the CBU, where the path integration memory is hypothesised to reside (***Stone et al., 2017***). Image adapted from the insectbraindb.org. This pseudo-colour image of the ant *Cataglyphis nodus* was created with original data from ***Habenstein et al. (2020)***. (**B**) Illustration of example outbound path and corresponding accumulated home vector. (**C**) Conceptual depiction of the path integration model with each of the eight columns encoding one of the cardinal directions in a Cartesian coordinate system (***Stone et al., 2017***). Rectangular boxes represent neuronal ensembles and shaded portions the activity level or stored value. The inputs are the current insect heading (population coding with eight cardinal directions around the animal, i.e. E-PG neurons ***Wolff and Rubin 2018***) and the current insect speed (encoded as the spike rate of TN neurons ***Stone et al. 2017***). Both the heading and speed neurons drive the eight speed integrating ‘memory units’. The inhibitory heading signals mask the speed signal, so the lower the heading neuron activity, the more the speed signal is integrated by the corresponding memory unit. The memory unit values form a sinusoidal pattern encoding the distance from the origin in its amplitude (*α*) and the direction to the origin in the horizontal location of the minimum.

In our model, each memory unit consists of a bump attractor encoding the agent’s displacement in the corresponding cardinal direction. We assumed that the bump attractors consist of neurons with a large enough time constant, resulting in a homing distance error increase rate comparable to that observed in ants (0.343 m^2^/h, see section Agent memory degradation model in Materials and Methods). During the waiting time period the bump attractors accumulated error at that rate. Then the agent was ‘released’ to return to its starting point (nest). As expected, the dispersion of the homing distance increased with the waiting time (***Figure 6A***i, and ***Figure 6A***ii). However, the median homing distance of the agent across trials did not significantly change with waiting time which is contrary to the monotonic decay observed in the animals (***Figure 6A***ii). Clearly, the animal’s memory follows dissipation dynamics that do not match the state degradation dynamics of bump attractor networks.

**Figure 6.**
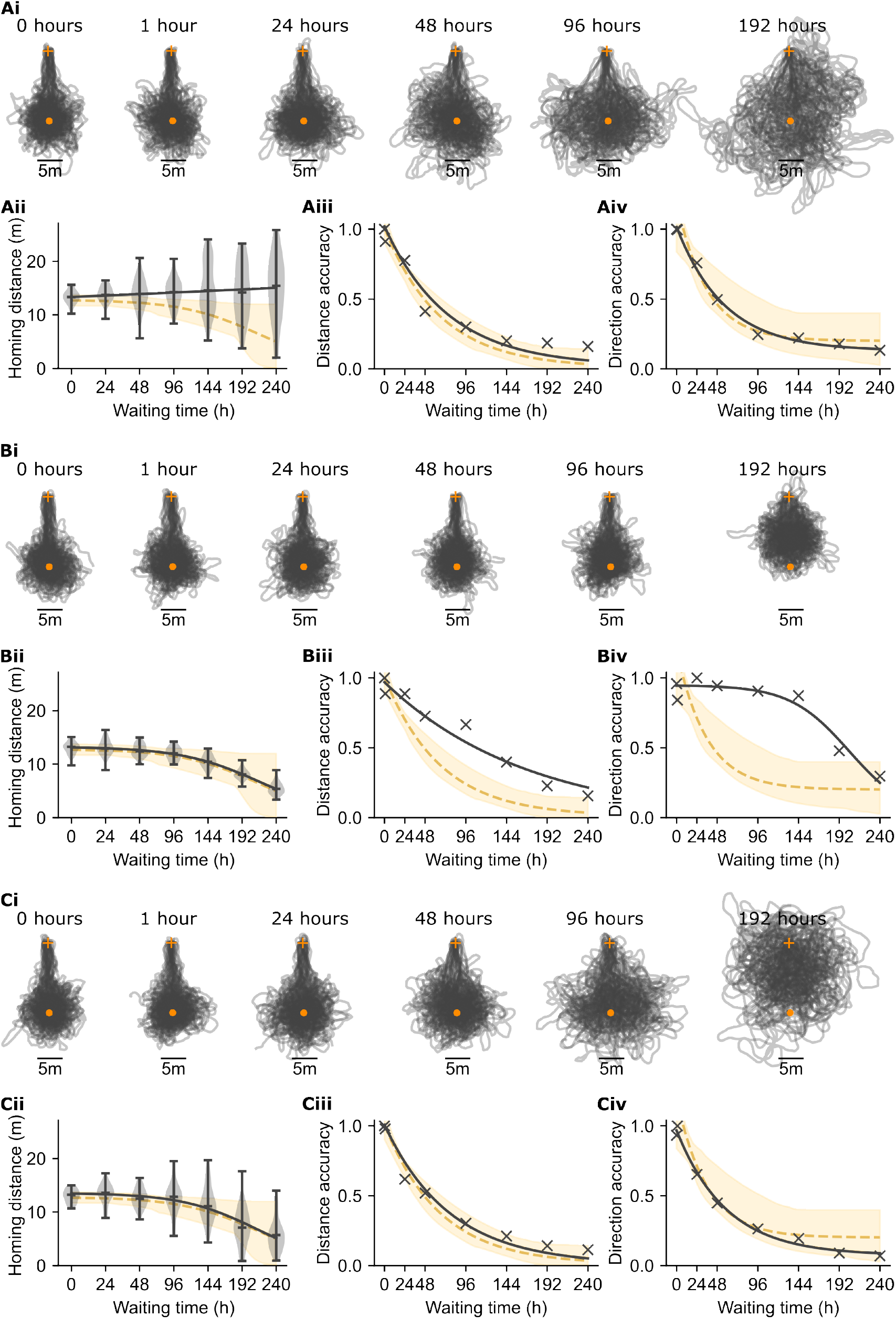
Comparisons of simulated agent homing to ant homing performance. Panels **A**, **B**, and **C** show the results obtained with three memory degradation models: (**A**) bump attractors exhibiting diffusive stochastic memory degradation; (**B**) time-dependent logistic memory decay (with parameters *K* = 1, *N*_*l*_ = 0.02, *r* = 0.018, see section Logistic decay model in Materials and Methods), and (**C**) the combined effect of time-dependent logistic decay and stochastic diffusive memory degradation (with parameters *K* = 1, *N*_*l*_ = 0.021, *r* = 0.021, and dispersion rate 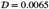, see section Hybrid logistic decay and diffusive degradation model in Materials and Methods). (**i**) Paths of an agent that was kept stationary (waiting) for different amounts of time before being released for homing. During the waiting time, the memory state accumulated error according to one of the three models (**A**, **B**, **C**). Then the agent is released to return to its origin as indicated by its home vector memory. The release location is indicated with an orange cross and the nest location with an orange disc. Once the agent reaches the location indicated by its memory, it searches following a looping trajectory resulting in the blob of overlapping paths centered at the expected home location. (**ii**) The homing distance (distance between the release point and the geometric centre of the search) as a function of the waiting time (the bars of the violin plots show the median, minimum and maximum). The black solid curve shows the regressed logistic function to the simulation data. The yellow dashed line shows the regressed logistic function to the ant homing distance data and the yellow shaded region indicates the 95% confidence intervals of the function fits as in ***Figure 4B***. (**iii**) The homing distance accuracy as a function of the waiting time (the ordinate shows MAD^−1^ in *m*^−1^ units). The exponential function (yellow dashed curve) *y* = *e*^(*bx*)^ + *c* was regressed to the ant homing data from ***Ziegler and Wehner*** (***1997***) with parameters *b* = −0.015 h^−1^, and *c* = 0.009 m^−1^. The yellow shaded region indicates the 95% confidence intervals for the function fits. The crosses (x’s) mark the medians of model simulation results and the black curve the exponential function regressed to the simulation data points with parameters *b* = −0.013, *c* = 0.020 (R^2^=0.94) for model **A**, *b* = −0.006, *c* = −0.030 (R^2^=0.95) for model **B**, and *b* = −0.013, *c* = 0.008 (R^2^=0.97) for model **C**. (**iv**) The homing direction accuracy as a function of the waiting time (the ordinate shows *σ*^−1^ in degree^−1^ units). The exponential function (yellow dashed curve) *y* = *e*^(*bx*)^ + *c* was regressed to the ant data from ***Ziegler and Wehner*** (***1997***) with parameters *b* = −0.028 h^−1^, and *c* = 0.201degree^−1^. The yellow shaded region indicates the 95% confidence intervals of the function fits. The crosses (x’s) mark the model simulation medians and the black curve is the exponential function regressed to the simulation data points with parameters *b* = −0.018, *c* = 0.130 (R^2^=0.99) for model **A**, no parameters were found for model **B** (the curve shown in panel **Biv** is the logistic function with parameters *K* = 0.948, *N*_*l*_ = 0.003, *r* = 0.028, resulting in R^2^=0.94), and *b* = −0.017, *c* = 0.073 (R^2^=0.99) for model **C**.

We next investigated the agent homing performance when using an alternative memory degradation model. In this alternative model, the memory decayed over time following the logistic function observed in ants (***Figure 4B***), instead of the diffusive stochastic degradation produced by the bump attractor memory model. With this memory decay model the homing distance decreased monotonically as expected, but its dispersion remained constant (see the violin plots in ***Figure 6B***ii), instead of increasing with the waiting time (as in ***Figure 6A***ii). In addition, the directional accuracy was not compatible with ant performance (***Figure 6B***iv). Therefore, the pure logistic memory decay is not a sufficient model of the animal memory degradation.

Finally, we tested a hybrid model, combining the logistic memory decay with diffusive stochastic state loss. With this hybrid model the agent’s homing behaviour indeed resembled that of the ants (***Figure 6C***). The homing distance decayed with the waiting time, the dispersion increased, while both the distance and direction accuracy were within the confidence intervals of animal behaviour (***Figure 6C***ii–iv). Therefore, the hybrid model was the only one that reproduced all the measures of ant homing behaviour.

## Discussion

We investigated the plausibility of the bump attractor model as the mechanism underlying path integration memory in ants. To this end, we compared the temporal dynamics of the path integration memory of *Cataglyphis fortis* ants with those of bump attractors encoding the agent’s displacement in respect to its starting point. We tested the bump attractor hypothesis with respect to the temporal dynamics of increase in the homing distance error, the number of required neurons, and the required neuronal time constants.

The bump attractor hypothesis predicts that the error in the ants’ homing distance would increase as a function of the time intervening before release. This is indeed the case in *Cataglyphis fortis* ants. The bump attractor hypothesis also predicts that the bump attractor network must consist of a considerable number of neurons or of neurons with adequately high time constants to be stable enough at the ecologically relevant time scale. However, the corporeal reality of the insect brain renders this last prediction inadmissible since it neither contains enough neurons nor are they known to have the high time constants required. But even in the case that these conditions are satisfied by a yet unobserved physiological property substantially increasing the neuronal time constant, we showed that the state value drift dynamics predicted by the bump attractor hypothesis do not match the observed dynamics of the animal’s homing distance. All our attempts to coax the bump attractor into reproducing the animal homing distance dynamics have failed. We then investigated alternative models of memory degradation and found that the underlying memory mechanism must be subject to the combined effect of logistic decay and diffusive stochastic noise to reproduce behaviour resembling that of the animals.

In the present work, we assumed that the homing distance degradation in ants is due to a degradation in the animal’s home vector memory. It is not inconceivable that other factors might be involved in the observed behaviour, such as the motivational state of the animals. However, there was no evidence of such an effect in the experiments since the animals continued searching for their nest, indicating that they were motivated to return to it. We also assumed a linear relationship between memory and homing distance. This relationship might be more complex and further research combining behavioural with physiological observations might be required to elucidate its exact nature.

A further consideration is that the biophysical parameters used in our models are based on measurements performed on neurons *in vitro*, typically at room temperatures. The desert ants, *Cataglyphis fortis*, are active at much higher temperatures which unavoidably affect the biophysical properties of their neurons. At higher temperatures, the membrane ion-channels become faster (***Frankenhaeuser and Moore, 1963***; ***Tang et al., 2010***), their conductance increases (***Volgushev et al., 2000***), and the synaptic transmission is faster (***Postlethwaite et al., 2007***). The structural properties of the membrane are also affected, resulting in decreased capacitance and time constant (***Volgushev et al., 2000***), while the neuronal channel noise level also decreases (***Faisal et al., 2005***, ***2008***). The exact effect of the higher temperature on the neurons is complex and further investigation is needed. However, the magnitude and polarity of the effects of temperature seem unlikely to result in a significant coordinated improvement of the bump attractor’s stability.

Our study is limited to ants, however the central complex — the brain area where the path integration circuitry is believed to be located — is conserved across species (***Heinze et al., 2013***; ***Pfeiffer and Homberg, 2014***; ***Seelig and Jayaraman, 2015***; ***Weir and Dickinson, 2015***; ***Heinze, 2015***; ***Turner-Evans and Jayaraman, 2016***; ***el Jundi et al., 2018***; ***Franconville et al., 2018***). Therefore, our results are likely to extend to other insect species. However, justifying such a generalisation would require further studies of homing accuracy in other insect species.

Despite our attempts to address every one of the model’s limitations, the bump attractor hypothesis failed to withstand all our tests. Memory models with slow dynamics have been explored in previous work (***Lisman and Goldring, 1988***; ***Koulakov et al., 2002***; ***Fransén et al., 2006***; ***Nikitchenko and Koulakov, 2008***; ***Mongillo et al., 2008***); however, the bump attractor model is not only limited by the duration of state maintenance but also the dynamics of state degradation. We illustrated, however, that a memory degradation model combining the diffusive state degradation with a monotonic state decay following the logistic function can reproduce the animal homing behaviour.

Our work casts doubt on the bump attractor memory hypothesis and rekindles the quest for the memory substrate of path integration. The presented findings provide important bounds to the memory degradation dynamics and therefore the range of plausible biophysical processes that could underlie the path integration memory.

## Materials and Methods

### Data extraction and processing

The ant behaviour data were extracted from ***Ziegler and Wehner*** (***1997***) and ***Cheng et al**.* (***2005***). The extracted data points as well as the data produced from simulations in this paper were fitted using the curve_fit function of SciPy’s optimize Python package with curve parametrisation as described in ***Figure 2***, ***Figure 3***, ***Figure 4***, and ***Figure 6***. The slopes of the curves in ***Figure 2A*** and ***Figure 3A*** were calculated with linear regression on the data points.

The rate of increase in the variance of the bump’s expected location is inversely proportional to the number of neurons in the attractor network *N* (***Burak and Fiete, 2012***). Therefore, the rate of increase of MAD^2^ (squared median absolute deviation) would also be inversely proportional to the number of neurons *N*. We use MAD^2^ per time unit as the measure of dispersion rate, 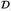, in m^2^ s^−1^ units. To extrapolate from the simulation-derived data points to higher values of *N*, in ***Figure 2D***, we used linear regression to the logarithm transformed data points

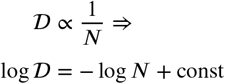

where 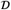 is the dispersion rate measured as squared median absolute deviation per second in m^2^ s^−1^ units.

Theoretical work suggests that the rate of increase in the variance of the bump’s expected location is bounded from below by a function inversely proportional to 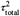, where *τ*_total_ is the sum of the time constants in the attractor network (***Burak and Fiete, 2012***). In ***Figure 3B***, the function 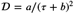 was regressed to the dispersion rate 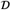 vs. the membrane time constant *τ* used in the simulations.

The confidence intervals in ***Figure 2B***, ***Figure 2D***, ***Figure 3B***, ***Figure 4B***, and ***Figure 6*** were estimated using bootstrapping.

### Bump attractor network model

The bump attractor network model implementation used in our experiments was derived from the code of ***Gerstner et al**.* (***2014***). Our source code is available at https://github.com/johnpi/Pisokas_Hennig_2022_Attractor_Based_Memory. To avoid the effect of boundary conditions at the edges of a line attractor, we used a ring attractor network topology and we made sure our experiments were confined to activity bump shifts smaller than ±180°. In this way, for the purposes of our experiments the topology of the circuit was indistinguishable from an actual line attractor.

The bump attractor network model used in the experiments had uniform global inhibition and structured lateral excitatory synapses with efficacies that followed a Gaussian profile. The skewed Gaussian formulation described below was used for determining the synaptic efficacies and allowed us to choose between Gaussian and skewed Gaussian profiles. The excitatory synaptic efficacies *w*(*x*_*i*_, *x*_*j*_) were therefore determined by the following formulation. Let *ϕ*(*x*) denote the Gaussian probability density function

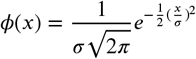

where *σ*^2^ = 20 is the variance (width) of the Gaussian profile. Let ϕ(*x*) denote the cumulative distribution function of the Gaussian

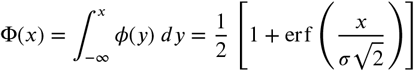

where *er f*(·) is the error function. Then the probability density function of the skewed Gaussian and thus the synaptic weights are given by

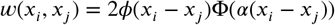

where *α* is the skewness factor, with *α* = 0 resulting in the Gaussian probability density function. In the case of simulations with systematically biased bump drift we used *α* = −0.0005 (***Figure 4***). *x*_*i*_ and *x*_*j*_ are the positions of the presynaptic and postsynaptic neurons, respectively, and *w*(*x*_*i*_, *x*_*j*_) ∈ [−1, 1] is the synaptic connection weight from neuron *x*_*i*_ to neuron *x*_*j*_. The values of *x*_*i*_ and *x*_*j*_ are periodic integers in the range [0, *N* − 1], where *N* is the number of excitatory neuronal units in the circuit and the difference *x*_*i*_ − *x*_*j*_ is calculated by modulo *N* subtraction, as in ***Brody et al. (2003)***. The values of *w*(*x*_*i*_, *x*_*j*_) determined the synaptic efficacy profile of the excitatory synapses while the inhibitory synapses had uniform efficacy.

The encoded distance values were in the range [0, 100]. This interval was mapped to the excitatory neurons around the ring attractor. Each of the excitatory neurons around the ring attractor was tuned to a distance value in the series 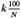, for *k* ∈ 0, 1, 2, … , *N* − 1. The bump’s location was decoded as a population vector for the purposes of the analysis presented here.

### Neuron model

We used two neuronal models in our experiments. The first one, the base model, utilised leaky integrate and fire neurons with AMPA, NMDA and GABA_A_ receptor channels. The inclusion of NMDA receptors endows the network with regular activity, since a network with only AMPA and GABA_A_ receptors produces synchronous spiking activity (***Compte et al., 2000***).

The second neuron model, utilised the neurotransmitter receptors of the first model and in addition mACh (muscarinic acetylcholine) receptors that metabotropically opened clusters of cooperative non-specific cation channels that provided the graded persistent spiking observed in the entorhinal cortex layer V neurons (***Egorov et al., 2002***, ***2006***).

The model parameter values were set to values consistent with evidence from measurements in *D. melanogaster* and other species when available (***Gouwens and Wilson, 2009***; ***Rohrbough and Broadie, 2002***; ***Sheeba et al., 2008***). The intrinsic neuronal properties for the excitatory neurons of the base model were set to a membrane capacitance *C*_*m*_ = 2nF, leak conductance *g*_*L*_ = 100 nS, reversal potential *E*_*L*_ = −70 mV, threshold potential *V*_*th*_ = −50 mV, reset potential *V*_*res*_ = −60 mV, and absolute refractory period *τ*_*ref*_ = 2ms. For the inhibitory neurons of the base model the corresponding values were *C*_*m*_ = 2nF, *g*_*L*_ = 200 nS, *E*_*L*_ = −70 mV, *V*_*th*_ = −50 mV, *V*_*res*_ = −60 mV, and *τ*_*ref*_ = 1ms.

In the second model, the intrinsic neuronal properties for the excitatory neurons with the non-specific cation conductance were set to a membrane capacitance *C*_*m*_ = 2nF, leak conductance *g*_*L*_ = 2nS, reversal potential *E*_*L*_ = −63 mV, threshold potential *V*_*th*_ = 10 mV, reset potential *V*_*res*_ = −60 mV, and absolute refractory period *τ*_*ref*_ = 3ms. For the inhibitory neurons of the second model the values were *C*_*m*_ = 2nF, *g*_*L*_ = 2nS, *E*_*L*_ = −63 mV, *V*_*th*_ = 10 mV, *V*_*res*_ = −60 mV, and *τ*_*ref*_ = 2ms.

For modifying the membrane time constant of the neurons the membrane capacitance was adjusted accordingly using the equation

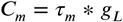

where *τ*_*m*_ is the desired time constant. The external input that set the initial bump location was provided by neurons forming synaptic connections with the excitatory neurons that were mediated by AMPA receptors, as in ***Compte et al. (2000)***. The maximum conductance of the AMPA receptor channels for the excitatory cells was *G*_*ext*,*E*_ = 3.1 nS and for the inhibitory cells *G*_*ext*,*I*_ = 2.38 nS.

Postsynaptic currents were modelled as

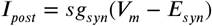

where *s* is the gating variable, *g*_*syn*_ the synaptic conductance, *V*_*m*_ the membrane potential, and *E*_*syn*_ the synaptic reversal potential (set to 0 mV for AMPA, NMDA, and mACh receptor channels and to −70 mV for GABA_A_ ones). The gating variables were modelled as in ***Compte et al. (2000)***. The decay time constants of the gating variables were set to 2 ms for AMPA, 10 ms for GABA_A_, 65 ms for NMDA, and 5 ms for mACh receptor channels.

The non-specific cation channels were modelled as clusters of cooperatively opening channels ***(Pfeiffer et al., 2020***). The cluster opening kinetics were governed by

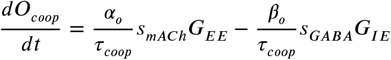

where *α*_*o*_ = 1 × 10^13^ and *β*_*o*_ = 5 × 10^13^ are the opening and closing rate factors, respectively. *τ*_*coop*_ = 100 s is the average cluster transition time constant. *s*_*mACh*_ and *s*_*GABA*_ are the channel gating variables. *G*_*EE*_ and *G*_*IE*_ are synaptic conductances described in ***Table 2***. The number of open clusters, *O*_*coop*_, was clipped to the range [0, 100]. The total cluster current was

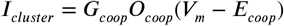

where *G*_*coop*_ = 7nS derived from the value used in ***Pfeiffer et al. (2020)*** and manually adjusted to obtain a functioning neuron model for the range of *O*_*coop*_ values. The synaptic conductances between neurons *i* and *j* were modelled as

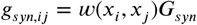

**Table 1.** Ant and bump attractor dynamics comparison.

| Parameter | Ants | Bump Attractor |
| --- | --- | --- |
| Distance degradation type | Monotonic | Random walk |
| Quickly resettable to zero state | Yes | Yes |
| Given $D = 0.343 \text{ m}^2/\text{h}$: | | |
| Number of neurons | thousands | millions |
| Time constant | <1 s | >8 h |

**Table 2.**
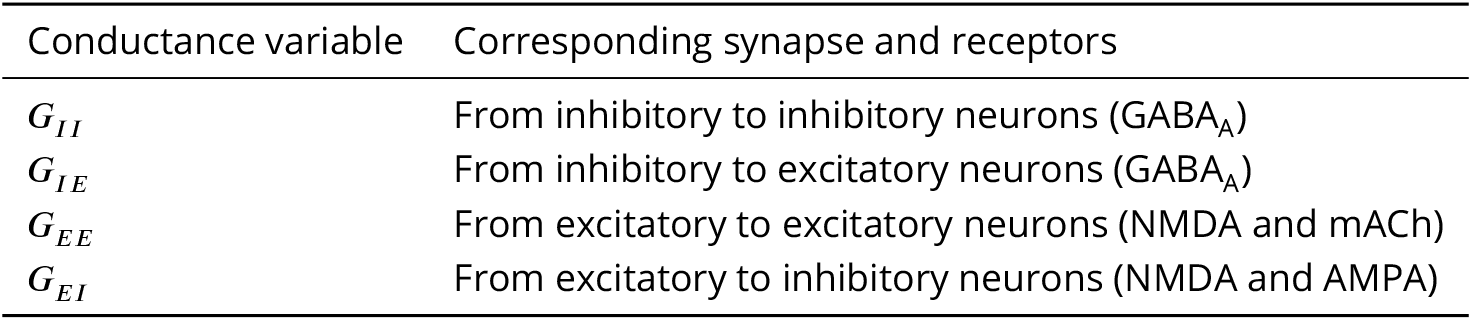
Synaptic conductances.

| Conductance variable | Corresponding synapse and receptors |
| --- | --- |
| $G_{II}$ | From inhibitory to inhibitory neurons (GABA<sub>A</sub>) |
| $G_{IE}$ | From inhibitory to excitatory neurons (GABA<sub>A</sub>) |
| $G_{EE}$ | From excitatory to excitatory neurons (NMDA and mACh) |
| $G_{EI}$ | From excitatory to inhibitory neurons (NMDA and AMPA) |

The synaptic conductances *G*_*syn*_ mediated by the AMPA, NMDA, GABA_A_, and mACh receptor channels were determined by following the procedure outlined in ***Brody et al. (2003)*** to obtain the formation of a stable activity bump. Subsequently, starting with the hand tuned values, the dual_annealing optimiser from SciPy’s optimize package was utilised to optimise the synaptic conductances using the objective function

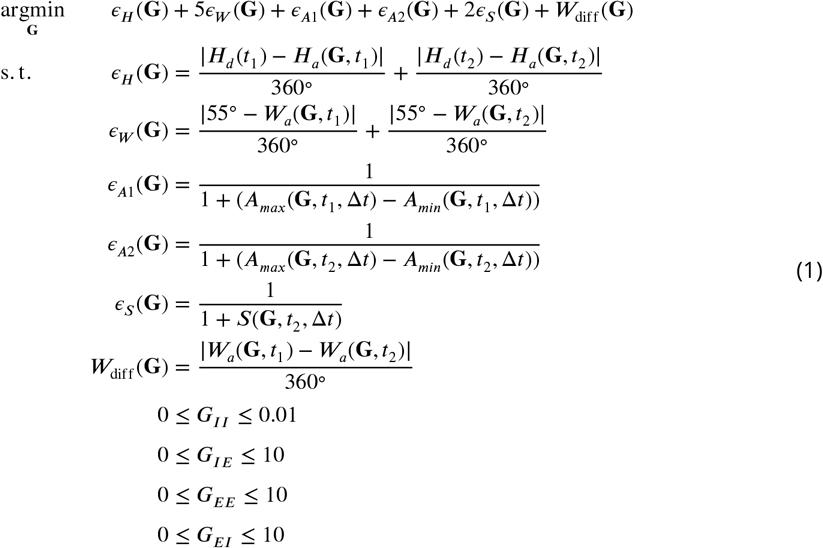

Where *ϵ*_*H*_ and *ϵ*_*W*_ are error factors measured as deviations from the desired values. *ϵ*_*A*1_, *ϵ*_*A*2_, *ϵ*_*S*_, and *W*_diff_ are factors penalising undesirable network behaviour. *H*_*d*_(*t*) is the desired activity ‘bump’ heading at time *t*, while *H*_*a*_(**G**, *t*) is the actual measured activity ‘bump’ heading at time *t* given a model with synaptic channel efficacies **G**. ***W***_*a*_(**G**, *t*) is the actual measured width of the activity ‘bump’ at time *t* (measured as the full width at half maximum). *A*_*max*_(**G**, *t*, Δ*t*) and *A*_*min*_(**G**, *t*, Δ*t*) are the maximum and minimum neuron spike rates across all excitatory neurons in the attractor network with channel conductances **G**, measured for a duration Δ*t* starting at time *t*. *S*(**G**, *t*, Δ*t*) ∈ [0, 1] is the proportion of the inhibitory neurons in the attractor network that have a spike rate greater than 30 impulses/s, measured for a duration Δ*t* starting at time *t*. *t*_1_ and *t*_2_ are sampling times located near the beginning and the end of the simulation, respectively. *ϵ*_*H*_(**G**) penalises deviation of the activity bump’s location from the desired location. *ϵ*_*W*_(**G**) penalises deviation of the activity bump width from the desired value of 55°. *ϵ*_*A*1_(**G**) and *ϵ*_*A*2_(**G**) reward the formation of a bump with larger amplitude. *ϵ*_*S*_(**G**) rewards increased spike rate activity across the inhibitory neuron population to avoid these neurons being completely silenced. *W*_diff_(**G**) penalises changes in the bump’s width between the beginning and the end of the simulation thus promoting sustenance of an activity bump. **G** is the vector [*G*_*II*_, *G*_*IE*_, *G*_*EE*_, *G*_*EI*_]. *G*_*II*_, *G*_*IE*_, *G*_*EE*_, and *G*_*EI*_ are the synaptic conductances as described in ***Table 2***. These were optimised separately for the base model and the model that included the non-specific cation conductances. For the base model the values were G_II_ = 0.3584 nS, G_IE_ = 0.4676 nS, G_EE_ = 0.133 35 nS, and G_EI_ = 0.122 64 nS for networks with 2048 neurons. The conductances were scaled according to the number of neurons used as in ***Brody et al. (2003)***. For the model that included the non-specific cation conductances the values were G_II_ = 0.0001 nS, G_IE_ = 0.005 nS, G_EE_ = 0.11 nS, G_EI_ = 1.339 624 49 nS, and the additional G_coop_ = 7.0 nS for networks with 2048 neurons. The constraint 0 ≤ *G*_*coop*_ ≤ 10 was added to the constraints of ***Equation 1*** when optimising for the last case.

### Neuronal noise

All neurons receive excitatory input modelled as uncorrelated Poisson spike trains with average spike rate as specified in the appropriate sections of the text. The neuronal noise used in ***Figure 2A***, ***Figure 2C***, ***Figure 2D***, ***figure Supplement 1***, ***Figure 4A***, ***Figure 4C***, and ***Figure 4D*** was modelled as Poisson noise with an average spike rate 5 impulses/s. This noise level was chosen because the attractor’s state dispersion rate reaches a plateau beyond Poisson neuronal noise with an average spike rate of 4 impulses/s (***Figure 2C***). Furthermore, this is a plausible spike rate near the lower end of background activity rates measured in neurons of the central complex (***Heinze and Homberg, 2007***, ***2009***). For ***Figure 3***, the Poisson neuronal noise was set to an average spike rate 1400 impulses/s to allow networks with higher neuronal time constants to sustain a bump of activity.

### Agent simulations

The agent simulations were based on the anatomically constrained model of ***Stone et al. (2017)***. The source code of the original work was modified and extended as described below and it is available at https://github.com/johnpi/Pisokas_Hennig_2022_Attractor_Based_Memory. The neurons were modelled as rate-based perceptrons with a sigmoid activation function. Independent Gaussian noise with *μ* = 0 and *σ*^2^ = 0.01 was added to the activation of each neuron and the resulting values were clipped to the interval [0, 1].

To reproduce agent paths resembling those of ants we added motor noise that is parametric to the home vector length, as in ***Pisokas et al. (2022)***. This motor noise was modelled as a Gaussian noise factor applied to the steering commands of the agent

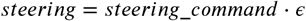

where *steering*_*command* is the steering command generated by the path integration model in homing mode, *steering* is the actual agent steering, and 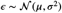 is a random variable sampled from the Gaussian distribution with *μ* = 1 and *σ* set to the motor noise level (***Equation 2***). The motor noise level was modelled as a decaying exponential function of the memory amplitude, as in ***Pisokas et al. (2022)***.

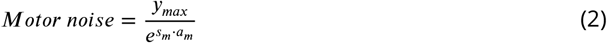

where *y*_*max*_ is the maximum motor noise level (corresponding to zero vector ant paths), and *s*_*m*_ is the slope of the exponential function. The memory amplitude *a*_*m*_ was defined as the difference between the maximum and minimum value along the sinusoidal pattern of memory unit values ***(Pisokas et al., 2022)***.

The translational velocity of the simulated agents during homing was iteratively updated using the formula

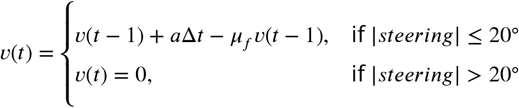

where *v*(*t*) is the agent’s velocity at time step *t*, *a* = 0.08 m/s^2^ is an acceleration constant, Δ*t* is the simulated time step duration, and *μ*_*f*_ = 0.15 is a friction constant. The velocity was reset to 0 m/s whenever the agent performed a turn larger than 20° simulating a stop on turning as observed in ants ***(Pisokas et al., 2022)***.

We performed a grid search following the procedure outlined in ***Pisokas et al. (2022)*** to find the combination of maximum motor noise level (*y*_*max*_ = 7) and slope (*s*_*m*_ = 9) of the decaying exponential that produces simulated paths that resemble the paths of both zero vector ants (zero memory amplitude) and full vector ants.

The inputs to the path integration model were the allocentric orientation of the agent and the speed of the agent’s motion (***Figure 5***, see also ***Stone et al. 2017***). Visual cues were not used in either the outbound or the homing part of the simulations; path integration was the sole navigation method utilised. The simulations had two stages, an outbound trip and a homing trip. For generating the outbound trip, the agent begun from a designated nest location and moved following a path generated by a filtered (smoothed) noise process, as previously described by ***Stone et al**.* (***2017***), until it reached a designated food location. Outbound agent simulations that did not result in the agent reaching the designated food location after 1500 simulation steps were disregarded. During the outbound trip, a home vector was accumulated that was encoded as the states of the eight memory units. Subsequently, an intervening waiting period was simulated during which stochastic memory state drift occurred, and then the simulation proceeded with the agent release in homing mode. The simulated homing paths were cut at 3000 steps from release allowing a sufficiently long observation period of the search behaviour to estimate the centre of search.

### Agent memory degradation model

#### Diffusive stochastic degradation model

In the agent behaviour simulations, for computational efficiency, the bump attractor memory dynamics were modeled with stochastic state diffusion. Therefore, the states of the memory units were updated by

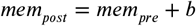

where *mem*_*pre*_ is the memory at the moment the agent was captured, *mem*_*post*_ the memory at the moment the agent was released, and

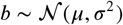

There were eight such memory unit variables one for each cardinal direction (***Figure 5C***). We set *μ* = 0 and *σ* to appropriate values corresponding to the effect of stochastic bump location drift for the different waiting periods. We set 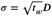, with *t*_*w*_ being the waiting period in hours and *D* determining the memory state dispersion rate. We experimentally determined that *D* = 28.8 produces the nearest approximation to the error increase rate observed in ant behaviour (dispersion rate 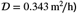).

#### Logistic decay model

For the logistic decay memory model, instead of using the diffusive stochastic degradation model, the memory values were multiplied by a time-dependent factor

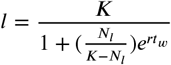

where *t*_*w*_ is the waiting time in hours, *K* = 1 sets the maximum value of *l* to 1, while *N*_*l*_ = 0.02 and *r* = 0.018 were determined using grid search to get the logistic function’s shape that minimizes the mean squared error between the agent homing distance and the ant homing distance over waiting time (***Figure 4B***). Note that these logistic function parameter values are different from the ones in ***Figure 4B*** because for the agent model, we regressed the logistic function to the normalised ant homing distance data (distance normalised to a maximum value of 1) so that *l* is in the interval [0,1]. The memory values were multiplied by the time-dependent factor *l* before release

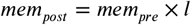

#### Hybrid logistic decay and diffusive degradation model

In the hybrid memory degradation model both the time-dependent logistic decay and diffusive stochastic degradation were applied on the memory values before release

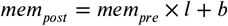

In this case the parameter values were *K* = 1, *N*_*l*_ = 0.021, and *r* = 0.021, while the dispersion rate determining factor was *D* = 0.0065. These parameter values were determined using grid search to minimize the sum of the mean squared errors between the agent simulation and the ant behaviour data for homing distance, homing distance error, homing distance accuracy, and homing direction accuracy.

## Acknowledgements

We would like to thank Barbara Webb for early discussion of the ideas developed in this paper and her comments on the draft. We would also like to thank Eve Marder and Stanley Heinze for early discussion of this work. Part of this work was funded by the Principal’s Career Development Scholarship, University of Edinburgh.

---

**Figure 3—figure supplement 1.**
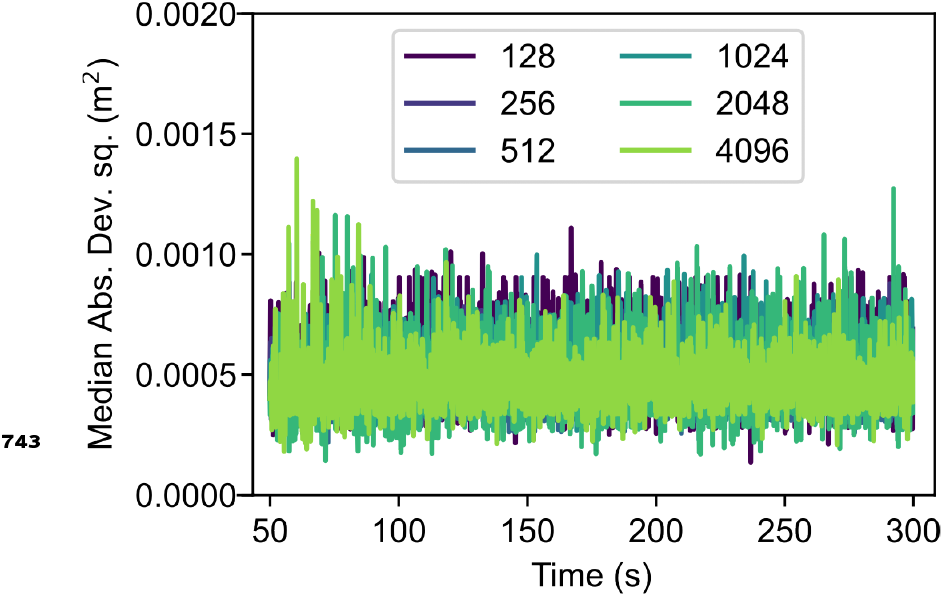
Bump attractor error increase using neurons with a persistent non-specific cation conductance. Simulation results show the error in the activity bump’s location over time for networks of different sizes. A homogeneous neuronal population was used. In these experiments the dispersion of the bump’s location was significantly smaller than in ***Figure 2A*** and ***Figure 3A***.

## Footnotes

1 The distance or the speed can be used for the calculation since the integral of speed over time gives the distance travelled.

